# The anti-obesity effect of fish oil in diet-induced obese mice occurs via the induction of heat production in brown but not white adipose tissue

**DOI:** 10.1101/2023.10.19.563064

**Authors:** Takahiko Obo, Hiroshi Hashiguchi, Eriko Matsuda, Shigeru Kawade, Kazuma Ogiso, Haruki Iwai, Koji Ataka, Osamu Yasuda, Aiko Arimura, Takahisa Deguchi, Akihiro Asakawa, Yoshihiko Nishio

## Abstract

**Aims/Introduction:** The ω3 polyunsaturated fatty acids in fish oil enhance heat production in adipocytes and exert anti-obesity effects, but the effects of fish oil on heat production in diet-induced obese (DIO) mice are unclear. In this study, we examined whether diets containing fish oil increased the expression of heat-producing genes in adipose tissue and increased body temperature in DIO mice, resulting in weight loss. We also examined fibroblast growth factor 21 (FGF21) levels in blood and the expression of the FGF21 gene in adipose tissue of DIO mice fed fish oil.

**Materials and Methods:** C57BL6/J mice were fed a lard-based high-fat diet for 8 weeks starting at 5 weeks of age and then divided into two groups: one group was fed a fish oil-based high-fat diet, and the other group was fed a lard-based high-fat diet continuously for another 8 weeks. Mice fed a control diet for 16 weeks from the age of 5 weeks served as the control group. Mice were dissected at 21 weeks and used for analysis.

**Results:** Mice fed a fish oil-based high-fat diet lost body weight gain, adipose tissue weight gain, and reduced insulin/leptin resistance. In addition, the rectal temperatures of mice fed a fish oil-based high-fat diet remained higher. The administration of fish oil increased the expression of heat-producing genes in brown adipose tissue (BAT) but did not alter heat-producing genes in inguinal white adipose tissue (WAT). In DIO mice fed a fish oil-based high-fat diet, the FGF21 expression in BAT increased. Furthermore, βklotho expression in BAT increased and the blood FGF21 concentration was decreased compared to mice fed a lard-based high-fat diet.

**Conclusions:** In DIO mice, fish oil was shown to increase rectal temperature and ameliorate obesity. Furthermore, fish oil enhanced heat production in BAT, but not WAT, in DIO mice.

## Introduction

Since 1980, the prevalence of obesity has doubled in more than 70 countries and continues to increase in most other countries [1]. In 2023, the World Obesity Coalition warned that more than half of the world’s population would be classified as obese or overweight [2]. The ω3 polyunsaturated fatty acids contained in fish oil (FO) are known to inhibit obesity in mice [3,4]. Beige adipose tissue, produced by browning of white adipose tissue (WAT), has a potent thermogenic capacity similar to that of brown adipocytes [5,6,7], and FO is observed to increase uncoupling protein 1 (UCP1) expression in both brown adipose tissue (BAT) and WAT [8], causing heat production and suppressing weight gain [4,9]. However, most of these studies on the thermogenic effects of FO in mice have been conducted in nonobese mice, and it is unclear whether FO exerts its thermogenic effects in obese mice. Okue et al. [10] reported that FO did not induce the expression of UCP1 in inguinal WAT in diet-induced obese (DIO) mice, but they did not report the effects of FO on BAT or body temperature. To determine whether FO has a thermogenic effect in obese animals, we fed FO to DIO mice and examined changes in body weight, rectal temperature, and expression of heat production-related genes in BAT and inguinal WAT.

FGF21 has been reported to contribute to thermogenesis by enhancing peroxisome proliferator-activated receptor-γ coactivator-1α (PGC1α) protein expression in adipose tissue and inducing UCP1 expression [11,12]. However, there are only a few reports on the expression of FGF21 in adipose tissue when FO is fed, and all studies have been conducted in nonobese mice [8,9]. It has been reported that FO administration increases FGF21 gene expression in both BAT and WAT in nonobese mice [9], but it is unclear whether similar changes are observed in DIO mice. Therefore, we also examined changes in the expression of the FGF21 gene in blood and adipose tissue using DIO mice fed FO.

## Methods

### Animals and diets

Four-week-old male C57BL/6J mice were purchased from KBT Oriental Co., Ltd. (Saga, Japan) and housed at 23 ± 1°C on a 12-h light/12-h dark cycle with ad libitum access to food and water.

After a 1-week acclimation period, mice were divided into two groups: one fed a control diet (CD) (D20051305M, Research Diets Inc., New Brunswick, USA) and the other fed a lard-based high-fat diet (LD) (D20051306M) for 8 weeks. The mice that were fed LD for 8 weeks were randomly assigned to one of the following regimens: continuation of a LD for another 8 weeks or introduced to a fish oil-based high-fat diet (FOD) (D20051307M) for another 8 weeks. The mice fed the CD were maintained on this diet for another 8 weeks (Fig 1). The CD contained 9% of calories from fat, 21% of calories from protein, and 70% of calories from carbohydrates (3.8 kcal/g), and the LD and the FOD contained 60% of calories from fat, 21% of calories from protein, and 19% of calories from carbohydrates (5.3 kcal/g) (Table 1). Food was provided to the mice every other day. To estimate daily food intake, the food weight of each day was subtracted from the initial food weight of the previous day. The mean calorie intake in each of the three groups was calculated using these data. Mice were weighed after a 6-hour fast at the end of each week of age. Rectal temperatures were measured weekly under feeding conditions at 7:00 a.m., which represented the end of the dark period. At 20 weeks of age, glucose and insulin tolerance tests (GTTs and ITTs, respectively) were performed on the mice, as described below. Because ITTs and GTTs were performed, rectal temperatures were measured under fasting conditions only at 20 weeks of age. At 21 weeks of age, the mice were sacrificed. All experimental procedures were reviewed and approved by the Laboratory Animal Committees of Kagoshima University Graduate School and were performed in accordance with the guidelines for the care and use of laboratory animals.

**Table 1.**
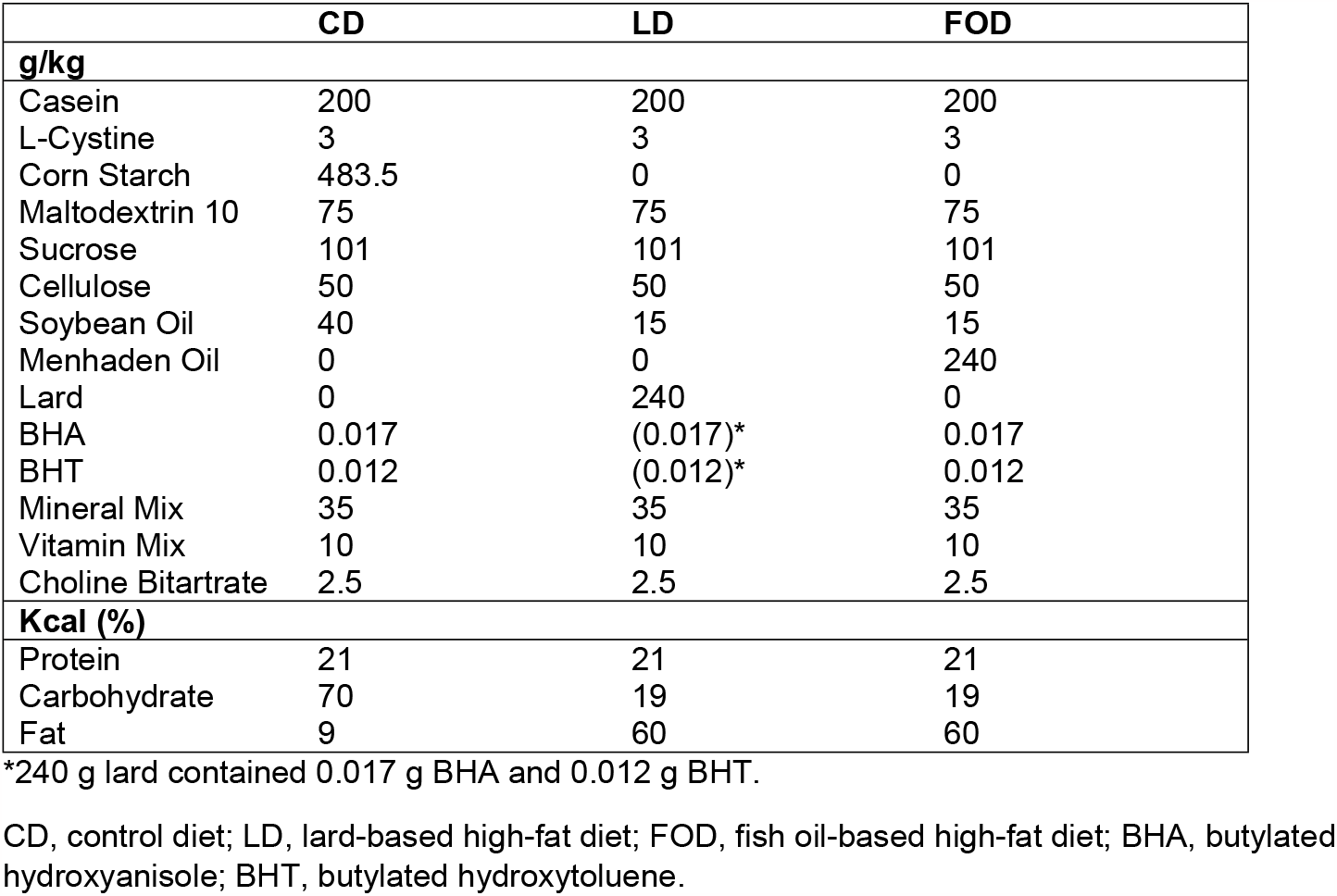
Composition of experimental diets.

|  | CD | LD | FOD |
| --- | --- | --- | --- |
| **g/kg** |  |  |  |
| Casein | 200 | 200 | 200 |
| L-Cystine | 3 | 3 | 3 |
| Corn Starch | 483.5 | 0 | 0 |
| Maltodextrin 10 | 75 | 75 | 75 |
| Sucrose | 101 | 101 | 101 |
| Cellulose | 50 | 50 | 50 |
| Soybean Oil | 40 | 15 | 15 |
| Menhaden Oil | 0 | 0 | 240 |
| Lard | 0 | 240 | 0 |
| BHA | 0.017 | (0.017)* | 0.017 |
| BHT | 0.012 | (0.012)* | 0.012 |
| Mineral Mix | 35 | 35 | 35 |
| Vitamin Mix | 10 | 10 | 10 |
| Choline Bitartrate | 2.5 | 2.5 | 2.5 |
| **Kcal (%)** |  |  |  |
| Protein | 21 | 21 | 21 |
| Carbohydrate | 70 | 19 | 19 |
| Fat | 9 | 60 | 60 |

\*240 g lard contained 0.017 g BHA and 0.012 g BHT.
CD, control diet; LD, lard-based high-fat diet; FOD, fish oil-based high-fat diet; BHA, butylated hydroxyanisole; BHT, butylated hydroxytoluene.

**Fig 1:**
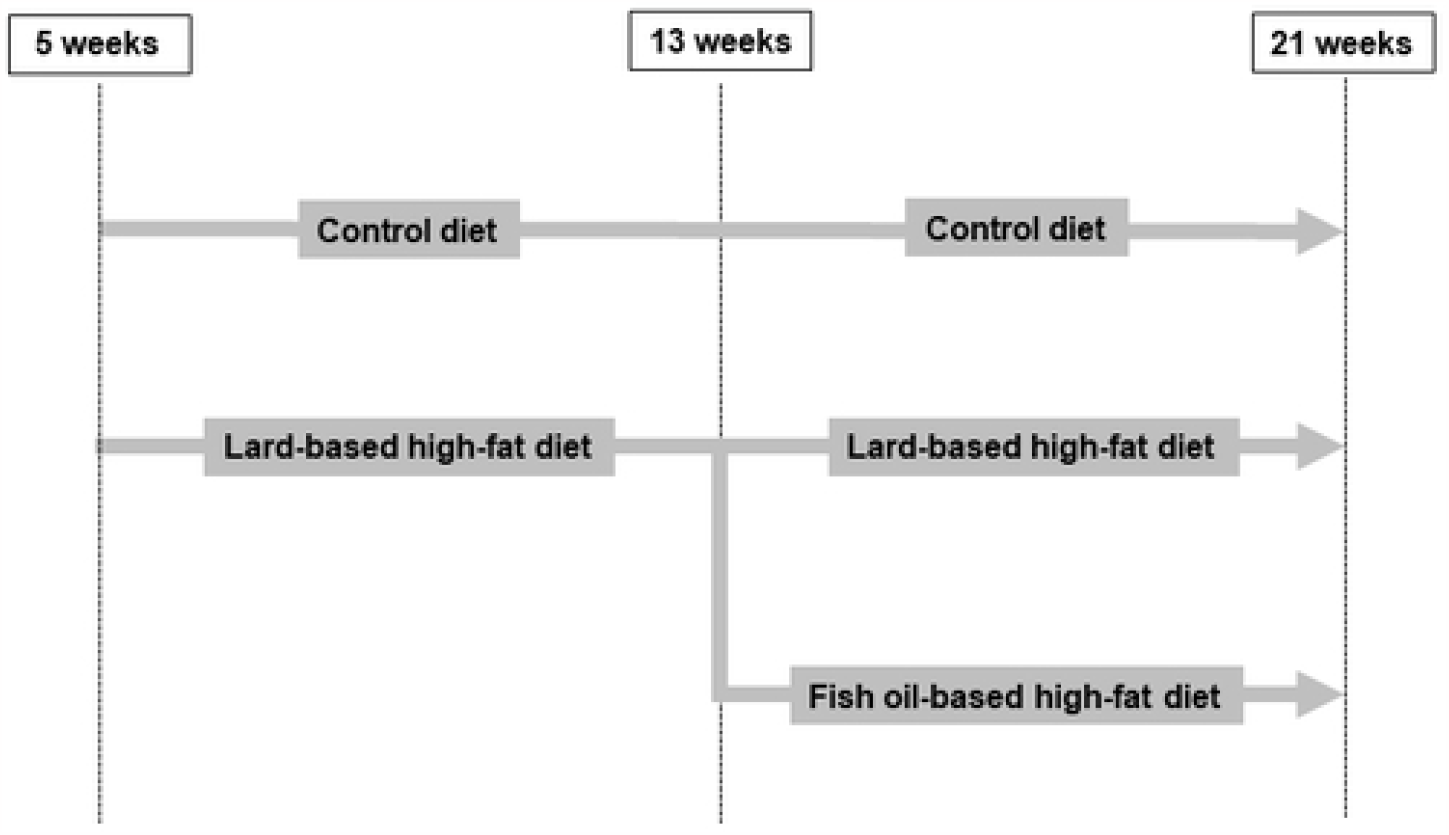
Experimental protocols. Five-week-old male C57BL/6J mice were fed a control diet (CD) and a lard-based high-fat diet (LD) for 8 weeks Mice fed a LD for 8 weeks were randomly divided into two groups: maintained for another 8 weeks on the LD or introduced to a fish oil-based high-fat diet (FOD) for another 8 weeks. The mice fed a CD were maintained on the CD for another 8 weeks. The CD contained 9% of calories from fat, 21% of calories from protein, and 70% of calories from carbohydrates (3.8 kcal/g), and the LD and the FOD contained 60% of calories from fat, 21% of calories from protein, and 19% of calories from carbohydrates (5.3 kcal/g) (Table 1). Food was provided to the mice every other day.

### Measurement of metabolic parameters

Blood was obtained from the mice, and plasma levels of nonesterified fatty acids (NEFAs), triglycerides (TG) and total cholesterol (TC) were measured by enzymatic colorimetry with LabAssay kits (Fujifilm Wako Pure Chemical Corporation, Osaka, Japan). Plasma FGF21 was measured using Mouse/Rat FGF21 Quantikine ELISA Kits (R&D Systems Inc., Minneapolis, USA). Plasma leptin was measured using a Mouse Leptin ELISA Kit (Proteintech Group Inc., Illinois, USA). Plasma insulin was measured using a mouse insulin ELISA kit (Morinaga Institute of Biological Science, Inc., Kanagawa, Japan). TG and TC were measured under fasting conditions. Because rectal temperature measurements were performed under nonfasting conditions, NEFAs, FGF21, and leptin were measured under nonfasting conditions for the purpose of investigating their relationship to body temperature.

### Glucose tolerance test and insulin tolerance test

The GTT was performed in the morning after an overnight fast. D-Glucose (2 mg/g body weight) was injected intraperitoneally. Capillary blood samples were collected using the tail cut method, and blood glucose was measured with Stat Strip XP3 (NIPRO Corporation, Osaka, Japan) before glucose injection and at 30, 60, and 120 min after glucose injection. The ITT was performed in the morning after an overnight fast. Insulin (Humulin R, Eli Lilly Japan K.K., Hyogo, Japan) was injected intraperitoneally at 0.1 mU/g body weight, and blood glucose was measured before insulin injection and at 30, 60, and 120 min after insulin injection.

### Histology

Twenty-one-week-old mice were intraperitoneally administered 90 mg/kg pentobarbital after inhaling anesthesia with isoflurane. The mice were systemically perfused with phosphate-buffered saline from the left ventricle and then perfused with 4% paraformaldehyde and fixed. After fixation, the livers were removed, and Oil red O staining was performed at the Bio-Pathology Laboratory (Oita, Japan).

### Quantitative real-time PCR and organ weights

Mice were anesthetized with pentobarbital 90 mg/kg intraperitoneally at 21 weeks of age, and inguinal adipose, brown adipose, epididymal adipose, mesenteric adipose, and liver tissues were harvested after general perfusion with phosphate-buffered saline. RNA was extracted from tissues with TRIzol Reagent (Life Technologies Japan Ltd., Tokyo, Japan) following the manufacturer’s instructions. Reverse transcription was performed by using High Capacity cDNA Reverse Transcription Kits (Thermo Fisher Scientific K.K., Tokyo, Japan). Quantitative real-time PCR was performed on a StepOnePlus Real-Time PCR system with TaqMan Fast Universal PCR Master Mix (Thermo Fisher Scientific K.K.) and the following gene-specific primers: Gapdh, Mm99999915_g1; Ucp1, Mm01244861_m1; Prdm16, Mm00712556_m1; Fgf21, Mm00840165_g1; βklotho, Mm00502002_m1; Adrb3, Mm02601819_g1; Ppargc1a, Mm01208835_m1; Ppara, Mm00440939_m1; Pparg, Mm01184322_m1; and Ffar4 Mm00725193_m1. Relative gene expression was calculated using the ΔΔCt method. The expression of target genes was normalized to that of GAPDH.

### Statistical analysis

All numerical values are expressed as the mean ± standard error of the mean. The two-sample t test was used for comparisons between the two groups. Comparisons between multiple groups were performed with one-way analysis of variance (ANOVA), and multiple comparison using the Tukey or Games-Howell post hoc analysis. Statistical significance was indicated by a *p* value < 0.05. All statistical analyses were performed using IBM SPSS Statistics 29.

## Results

### Feeding fish oil to DIO mice increased rectal temperature and improved obesity

As shown in Fig 2a, at 12 weeks of age, the DIO model mice (LD and FOD) weighed significantly more than the nonobese mice (33.44 ± 2.09 g vs. 27.16 ± 1.08 g, p < 0.001 by independent t test).

During the experimental period, the body weight of mice fed the LD increased in a time-dependent manner (Fig 2a, b). The body weight of mice fed the CD remained unchanged after 12 weeks. Mice fed the LD had a higher caloric intake per day than the other two groups(Fig 2c, d). Mice fed the FOD had a temporary decrease in caloric intake when switching from LD to FOD at 13 weeks, but thereafter, caloric intake was similar to that of mice fed CD. A temporary decrease in caloric intake following the change from LD to FOD has been reported in previous studies [13]. Rectal temperatures in the mice fed LD and FOD were similarly higher than those in mice fed the CD (Fig 2e, f). Thus, mice fed the LD showed higher caloric intake and higher rectal temperatures compared to mice fed the CD, whereas mice fed the FOD showed similar caloric intake to mice fed the CD but higher rectal temperatures, suggesting the mechanism of bodyweight reduction by FOD.

**Fig 2:**
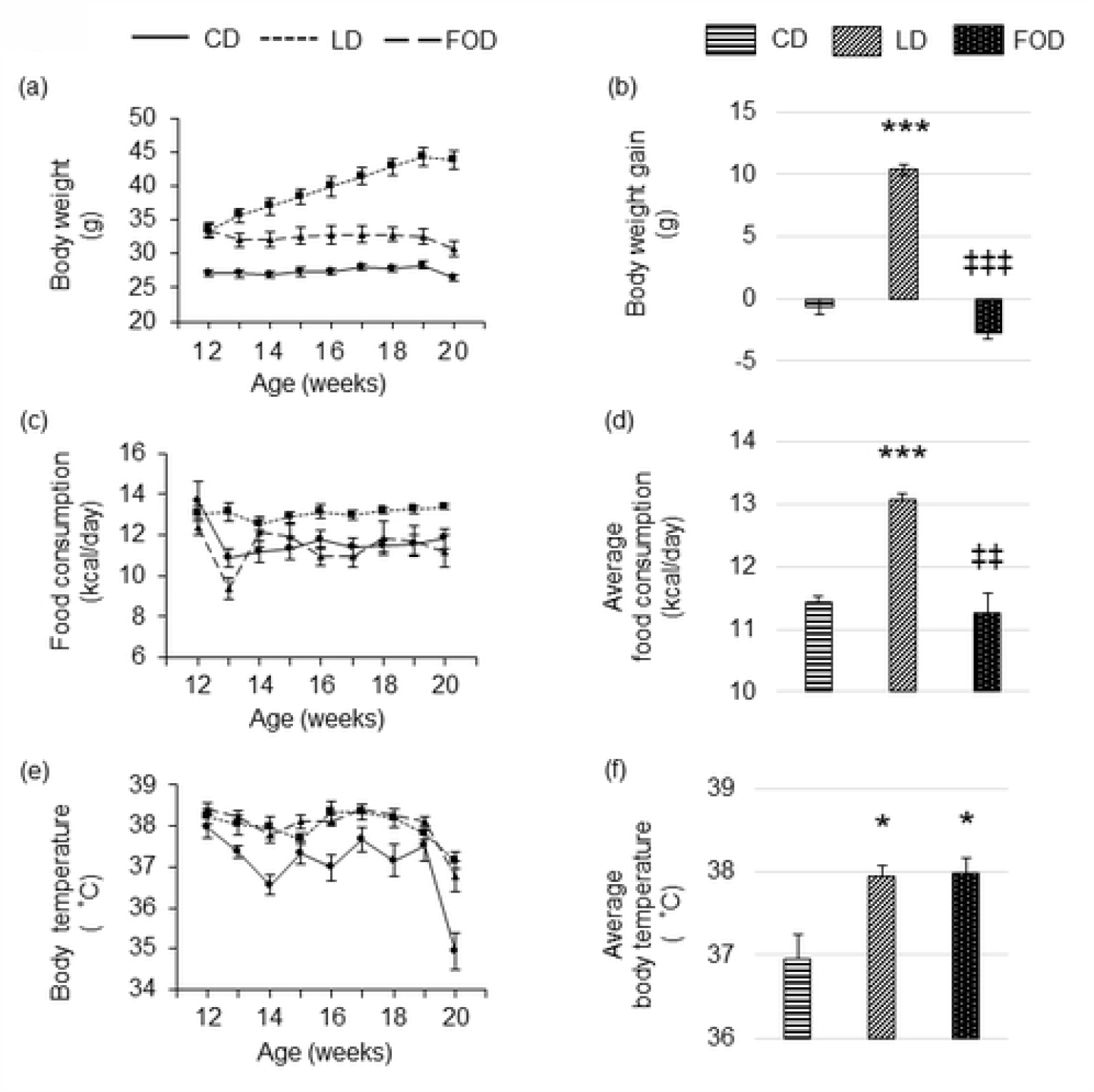
Fish oil increases rectal temperature and improves obesity. Feeding a FOD to DIO mice resulted in caloric intake similar to that of mice fed a CD, body temperature remained as high as that of mice fed the LD, and obesity improved. (a) Body weight; (b) body weight gain between 12 and 20 weeks of age; (c) food consumption; (d) average food consumption between 12 and 20 weeks of age; (e) body temperature; (f) average body temperature between 12 and 20 weeks of age. The data are presented as the mean ± standard error of the mean, *n* = 4-6 animals per group. * P < 0.05, *** P < 0.001 compared with CD group, ‡‡ P < 0.01, ‡‡‡ P < 0.001 compared with LD group using one-way ANOVA. FOD, fish oil-based high-fat diet; CD, control diet; LD, lard-based high-fat diet.

### Feeding fish oil to DIO model mice reduced subcutaneous and visceral fat and improved fatty liver

Consistent with body weight, WAT weights of the mesentery (Fig 3a), inguinal (Fig 3b), and epididymis regions (Fig 3c) were higher in mice fed the LD than in mice fed the CD, and mice fed FOD showed a significant reduction in weight in these tissues. Liver weight increased in both mice fed the LD and the FOD (Fig 3d), but Oil Red O-stained images of liver tissue revealed that remarkable fat deposition was observed only in mice fed the LD (Fig 3e), indicating that the increase in liver weight of mice fed the FOD does not reflect fatty liver. It has been reported that FO increases liver weight by inducing peroxisome proliferation [14].

**Fig 3:**
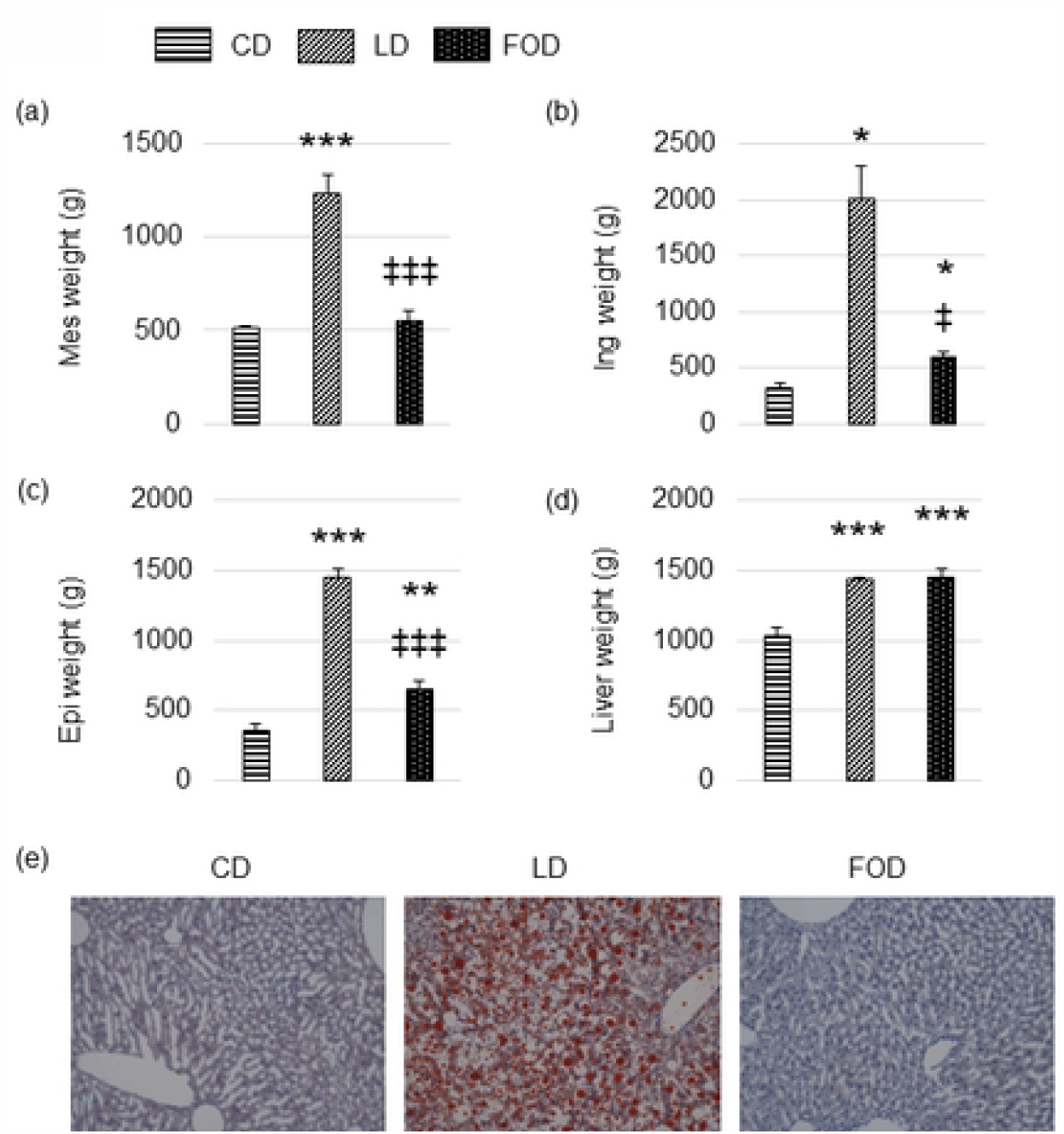
Fish oil reduces subcutaneous and visceral fat. Mice were dissected at 21 weeks, and liver and adipose tissue weights were compared. The weights of mesenteric fat, inguinal fat, and epididymal fat increased in LD-fed mice compared to CD-fed mice but were reduced in FOD-fed mice. Liver weights increased in both LD-fed and FOD-fed mice compared to CD-fed mice, but Oil Red O staining showed fat droplets only in LD-fed mice. (a) Mesenteric adipose tissue (Mes) mass; (b) inguinal adipose tissue (Ing) mass; (c) epididymal adipose tissue (Epi) mass; (d) liver mass; (e) representative Oil red o-stained liver histology. The data are presented as the mean ± standard error of the mean, *n* = 4-6 animals per group. * P < 0.05, ** P < 0.01, *** P < 0.001 compared with CD group, ‡ P < 0.05, ‡‡‡ P < 0.001 compared with LD group using one-way ANOVA. LD, lard-based high-fat diet; CD, control diet; FOD, fish oil-based high-fat diet.

### Feeding fish oil to DIO model mice improved glucose tolerance and insulin resistance

In the intraperitoneal ITT performed at 20 weeks, the area under the curve of blood glucose levels was larger in mice fed the LD than in mice fed the CD, and mice fed the FOD showed a significant reduction (Fig 4a, b). In the intraperitoneal GTT, the area under the curve of blood glucose levels in mice fed the LD was larger than that in mice fed the CD, and the mice fed the FOD showed a significant reduction (Fig 4c, d). Blood insulin levels were higher in mice fed the LD both before and after glucose administration (Fig 4e, f). This indicates that insulin resistance was improved by administering FO to DIO mice.

**Fig 4:**
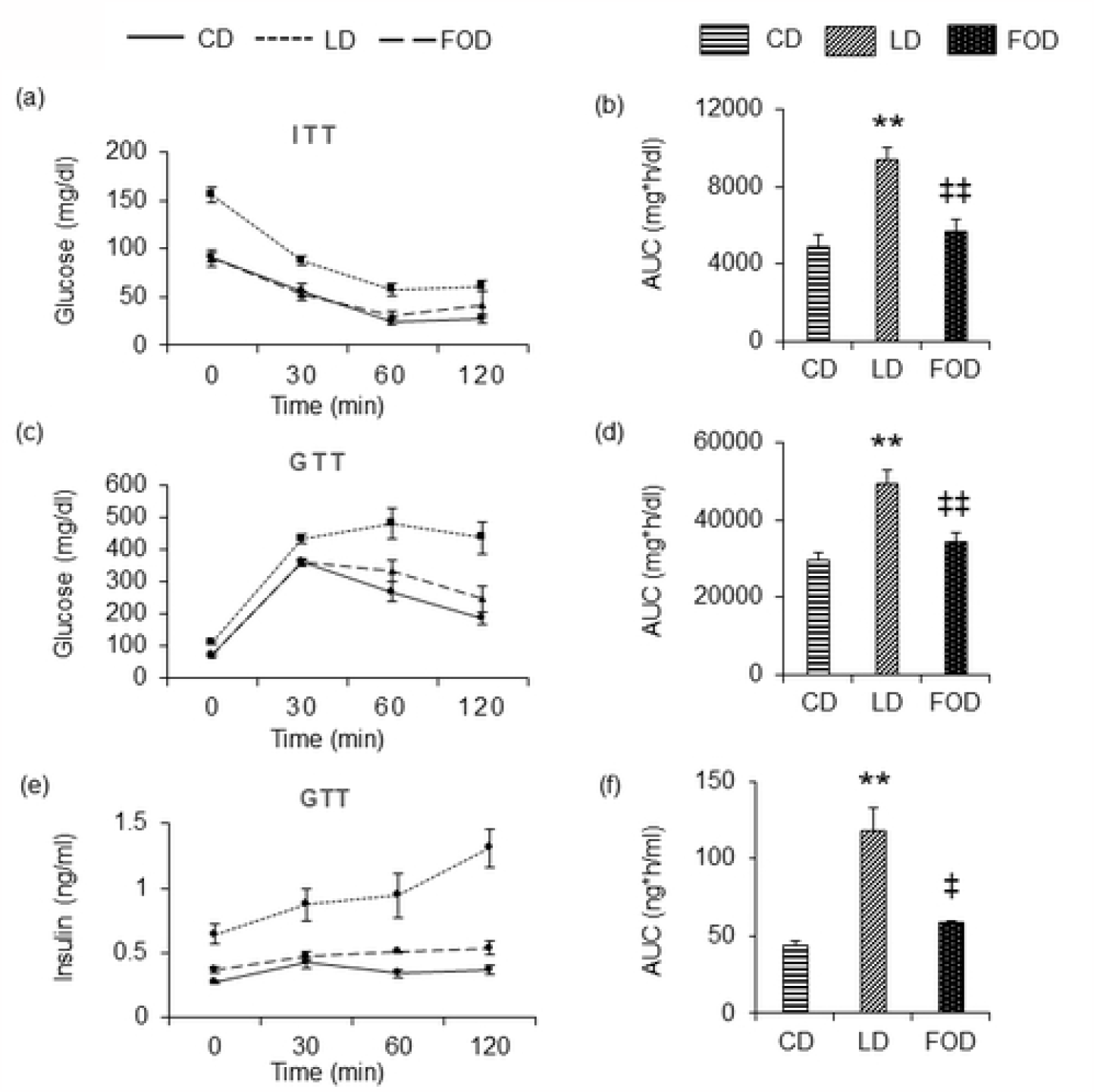
Fish oil improves glucose tolerance and insulin resistance. Mice fed the LD developed insulin resistance, which improved when mice were fed the with FOD. (a) Changes in blood glucose, as indicated by the ITT. (b) AUC of blood glucose levels during the ITT. (c) Changes in blood glucose, as indicated by the GTT. (d) AUC of blood glucose levels during the GTT. (e) Changes in plasma insulin, as indicated by the GTT. (f) AUC of plasma insulin levels during the GTT. The data are presented as the mean ± standard error of the mean, *n* = 4-8 animals per group. ** P < 0.01 compared with CD group, ‡ P < 0.05, ‡‡ P < 0.01 compared with LD group using one-way ANOVA. LD, lard-based high-fat diet; FOD, fish oil-based high-fat diet; ITT, insulin tolerance test; AUC, area under the curve; GTT, glucose tolerance test.

### Feeding fish oil to DIO mice decreased FBG, insulin, T-Chol, FGF21, and Leptin

Plasma total cholesterol (T-Chol) concentration was higher in mice fed the LD compared to mice fed the CD; furthermore, mice fed the FOD showed a significant reduction in T-Chol. Plasma triglyceride concentrations did not differ between groups. Blood free fatty acid concentrations tended to be lower in mice fed the FOD, but the difference was not significant. Blood FGF21 concentrations were significantly lower in mice fed the FOD than in mice fed the LD. Blood leptin levels were higher in mice fed the LD than in mice fed the CD, and mice fed the FOD showed a significant reduction, suggesting leptin resistance in mice fed the LD and an improvement when mice were fed the FOD (Table 2).

**Table 2.**
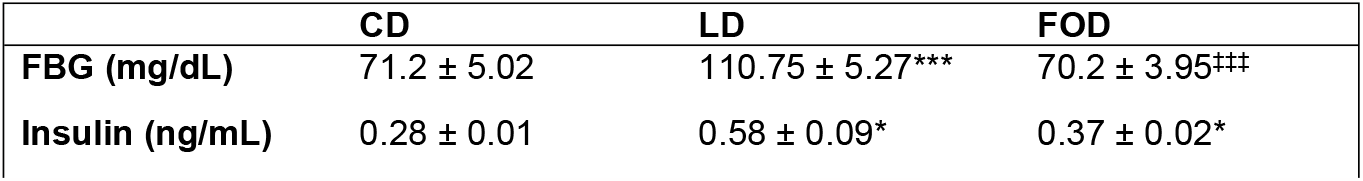
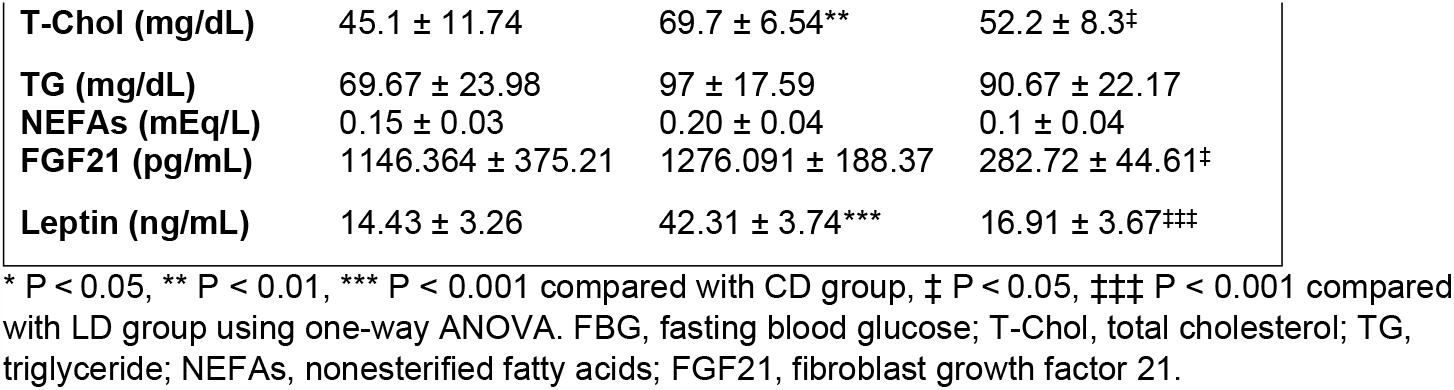
Plasma concentrations of various parameters in 21-week-old male C57BL/6J mice.

### Fish oil-fed and LD-fed DIO mice had increased expression of heat-producing genes in brown adipose tissue but not in white fat

To confirm the mechanism by which rectal temperature increased in mice fed LD and FOD, the mRNA expression of genes involved in heat production was measured in adipose tissue obtained from mice of each group (Fig 5, 6).

**Fig 5:**
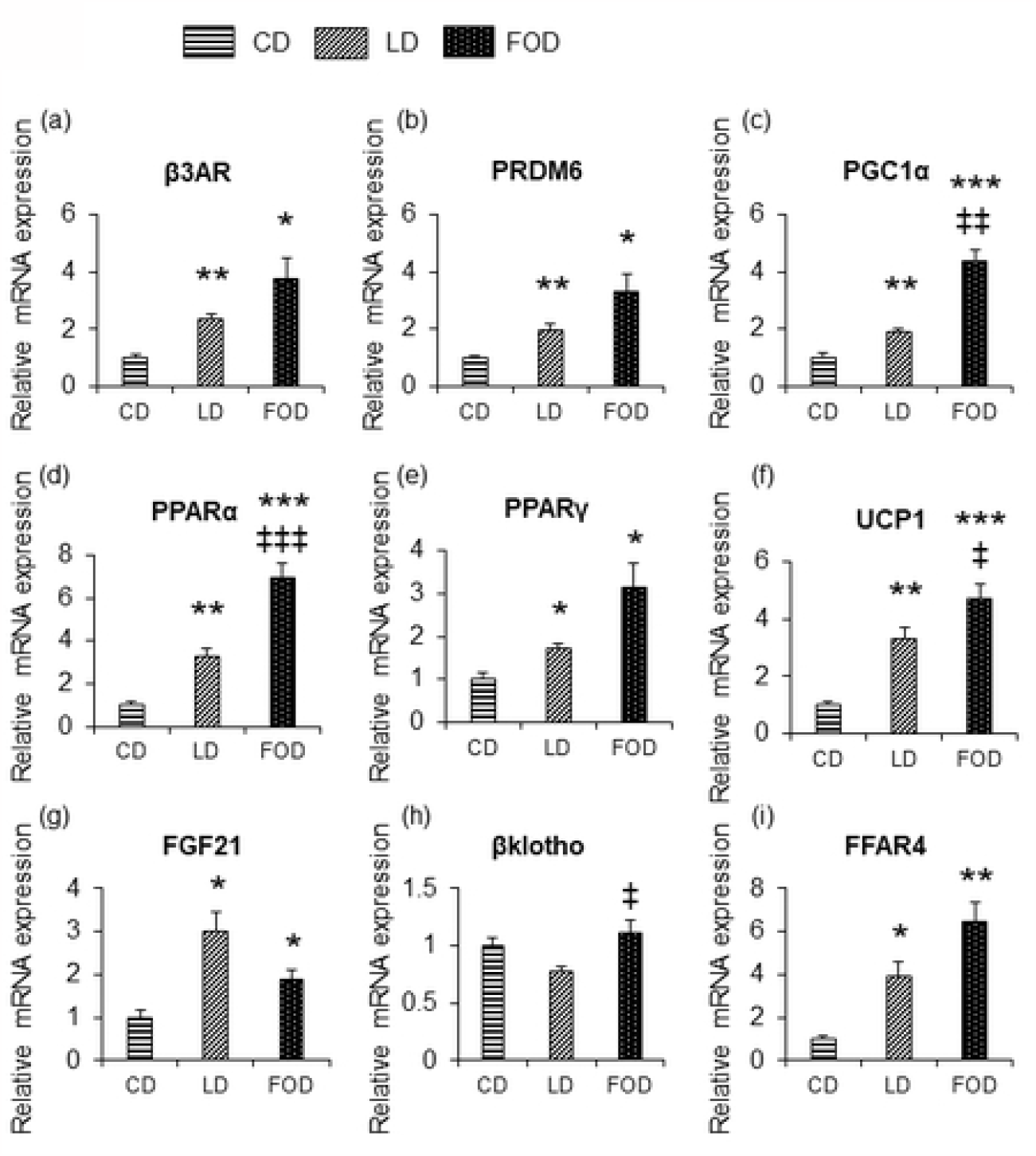
Fish oil increased the expression of heat-producing genes in BAT. The expression of genes involved in heat production increased in BAT of mice fed the LD and the FOD. Gene expression levels in BAT. (a) *β3AR*; (b) *PRDM16*; (c) *PGC1α*; (d) *PPARα*; (e) *PPARγ*; (f) *UCP1*; (g) *FGF21*; (h) *βklotho*; (i) *FFAR4*. The data are presented as the mean ± standard error of the mean, *n* = 6-7 animals per group. * P < 0.05, ** P < 0.01, *** P < 0.001 compared with CD group, ‡ P < 0.05, ‡‡ P < 0.01, ‡‡‡ P < 0.001 compared with LD group using one-way ANOVA. BAT, brown adipose tissue; *β3AR*, β3-adrenergic receptor; *PRDM16*, PR domain containing 16; *PGC1α*, peroxisome proliferator-activated receptor-γ coactivator-1α; *PPARα*, peroxisome proliferator-activated receptor-α; *PPARγ*, peroxisome proliferator-activated receptor-γ; *UCP1*, uncoupling protein 1; *FGF21*, fibroblast growth factor 21; *FFAR4*, free fatty acid receptor 4.

**Fig 6:**
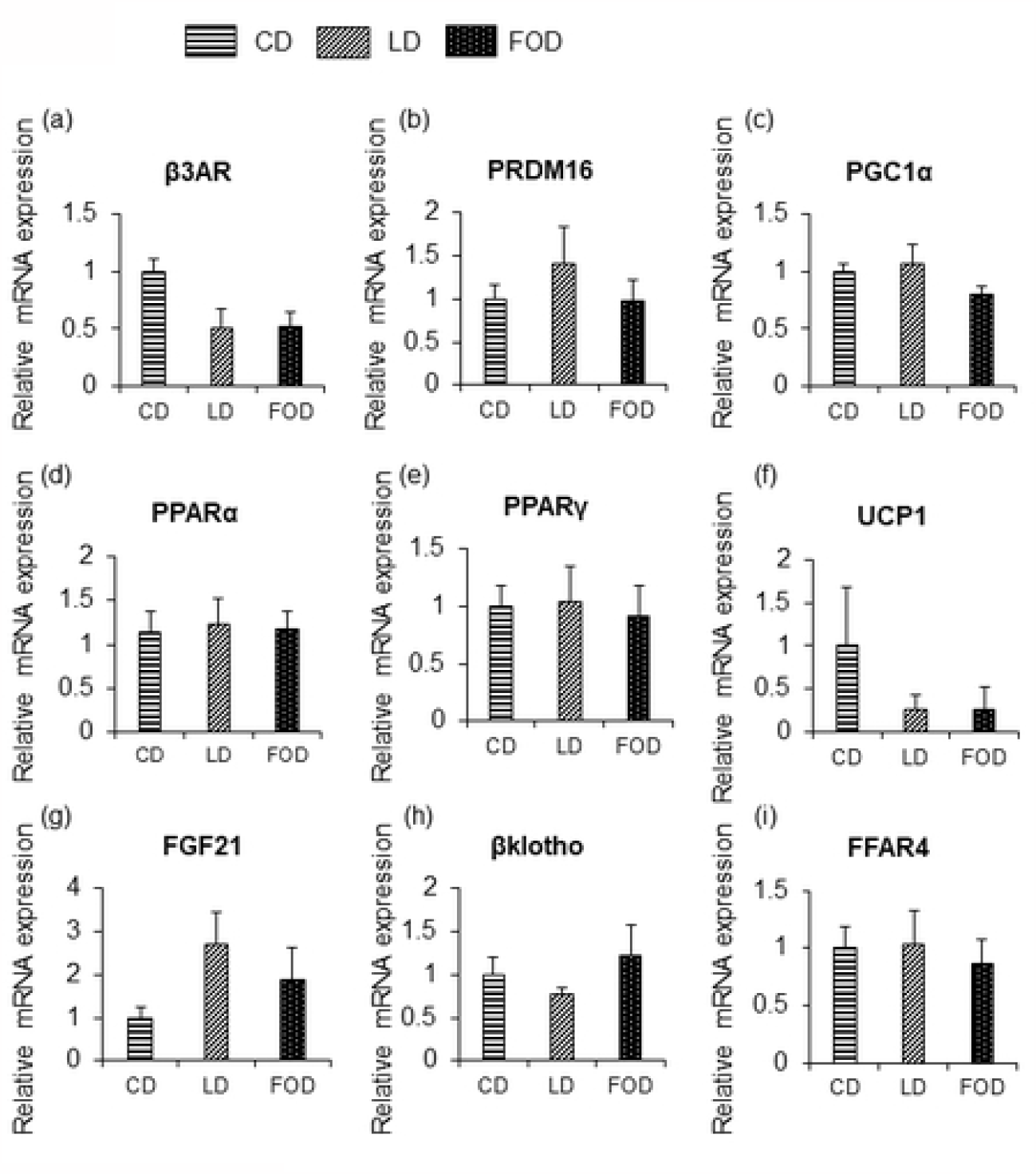
Fish oil did not increase the expression of heat-producing genes in inguinal WAT. No changes in gene expression were observed in inguinal WAT. (a) *β3AR*; (b) *PRDM16*; (c) *PGC1α*; (d) *PPARα*; (e) *PPARγ*; (f) *UCP1*; (g) *FGF21*; (h) *βklotho*; (i) *FFAR4*. The data are presented as the mean ± standard error of the mean, *n* = 6-7 animals per group. WAT, white adipose tissue; *β3AR*, β3-adrenergic receptor; *PRDM16*, PR domain containing 16; *PGC1α*, peroxisome proliferator-activated receptor-γ coactivator-1α; *PPARα*, peroxisome proliferator-activated receptor-α; *PPARγ*, peroxisome proliferator-activated receptor-γ; *UCP1*, uncoupling protein 1; *FGF21*, fibroblast growth factor 21; *FFAR4*, free fatty acid receptor 4.

FOD increased β3-adrenergic receptor (β3AR) mRNA expression by 3.7-fold in BAT (Fig 5a), PRDM16 by 3.3-fold (Fig 5b), PGC1α by 4.3-fold (Fig 5c), and UCP1 by 4.7-fold (Fig 5f) compared to mice fed the CD and increased PPARα (Fig 5d) and PPARγ (Fig 5e). Mice fed the LD also showed increased expression of genes involved in heat production, but not to the extent as observed in mice fed the FOD. The expression of the free fatty acid receptor FFAR4 increased in mice fed the LD and in mice fed the FOD compared to mice fed the CD (Fig 5i). FGF21 mRNA expression also increased in mice fed the LD and the FOD, showing a different trend from that of blood FGF21 levels (Fig 5g). βklotho, which acts as a receptor for FGF21, increased in mice fed the FOD compared to mice fed the LD (Fig 5h). Unlike the results related to BAT, no changes in gene expression involved in heat production were observed in inguinal WAT. (Fig 6a-i)

## Discussion

Feeding FO-containing diets to DIO mice reduced body weight, adipose tissue weight, and insulin and leptin resistance. In addition, the caloric intake of mice fed the FOD was reduced to the same level as that of mice fed the CD, while rectal temperature remained as high as that of mice fed the LD, thus resulting in weight loss. Administration of FO induced the expression of β3AR and increased the expression of heat-producing genes in BAT but did not alter heat-producing genes in inguinal WAT. These results indicate that FO acts as an anti-obesity agent via heat production in BAT, but not WAT, in DIO mice.

In this study, the rectal temperature of mice fed the FOD was similar to that of mice fed the LD and higher than that of mice fed the CD. In BAT and inguinal WAT, β3AR signaling induces PRDM16 and PGC1α via the β3AR-cAMP-PKA pathway and increases UCP1 expression in nonobese mice [6,15,16]. PPARα and PPARγ are also involved in heat production through this signaling pathway [15,17,18]. Consistent with these reports, in our study, the expression of β3AR and genes such as PRDM16, PGC1α, PPARα, PPARγ, and UCP-1 was increased in the BAT of mice fed the LD and the FOD.

DIO mice have increased heat production in BAT, due to the development of hyperleptinemia associated with obesity [4,19]. Leptin is produced and secreted by adipocytes and is responsible for energy homeostasis, and plasma leptin levels are observed to increase in obese animals [19,20]. Obese animals are resistant to leptin’s effect on appetite control but not to its centrally mediated sympathetic activating effects [19,21]. Blocking β3AR abolishes the increased thermogenic response in DIO mice [19]. On the other hand, mice fed a FOD did not develop hyperleptinemia in the present study, and FO is known to increase heat production in a leptin-independent manner [22]. It has been reported that FO increases β3AR expression in BAT and WAT via sympathetic activation [8], thereby causing increased expression of UCP1 and heat production in BAT and WAT in nonobese mice [9,23,24]. Since the induction of UCP1 mRNA expression in BAT and WAT by FO is abolished by beta-blocker administration [8], beta-receptor stimulation plays an important role in FO-induced heat production in adipose tissue. In the present study, we found that increased expression of genes involved in heat production in BAT occurred in DIO mice fed with FO. Although confirming elevated UCP1 mRNA expression does not definitively confirm that heat generation in BAT was activated, elevated rectal temperatures further suggest that heat production may have been activated. These results suggest that enhanced sympathetic signaling was involved in the increase in rectal temperature in both mice fed the LD and the FOD, but the mechanism of sympathetic activation in the two groups may be different.

In addition, the changes in gene expression involved in heat production identified in BAT of mice fed the LD and the FOD were not similarly observed in the inguinal WAT in this study. Okue et al. [10] reported that UCP-1 expression did not increase in the inguinal WAT of DIO mice fed diets containing FO for 4 weeks, which is consistent with our results. It has been reported that a high-fat diet decreases β3AR and UCP-1 expression in WAT [25,26]. Furthermore, Shin et al. [27] reported that in obese mice fed a high-fat diet, β3-adrenergic receptor agonists increased UCP1 gene and protein expression in BAT, while there was no increase in UCP1 gene and protein expression in WAT, suggesting WAT in obese animals fed a high-fat diet is less likely to be beige. In the present study, the expression of β3AR and UCP1 did not increase in inguinal WAT even after feeding FOD for 8 weeks resulted in an improvement in obesity, which suggests that the LD-induced reduction in β3AR signaling in WAT may persist long-term.

The results of this experiment suggest that the heat-producing effect of FO on obese individuals consuming a high-fat diet may depend on the function of BAT. In humans, the content of BAT, as detected by FDG-PET/CT, and activity is known to decrease with age [28]. Therefore, the anti-obesity effect of FO in obese individuals via heat production may diminish with age. Interestingly, meta-analyses of randomized controlled trials examining the weight-loss effects of FO in obese humans do not consistently demonstrate an associated weight-loss effect [29,30], but several studies in relatively young obese individuals up to age 40 have shown a weight-loss effect with FO [31,32]. Therefore, the effect of FO on weight loss through the promotion of heat production might be exerted only in young people with preserved BAT function.

FGF21 contributes to heat production by inducing UCP1 expression in adipose tissue [11,12], and it has been reported that FO feeding increases FGF21 expression in both BAT and WAT in nonobese mice [9]. However, feeding FO to DIO mice in the present study increased FGF21 expression only in BAT and not in inguinal WAT. Since β3 adrenergic receptor signaling is known to increase FGF21 mRNA expression in BAT without an increase in blood FGF21 levels [33,34], there may be a link between the observation that FO feeding increased β3AR mRNA expression only in BAT and not in WAT of DIO mice and that FGF21 expression increased only in BAT. The fatty acid receptor FFAR4 is highly expressed in adipose tissue [35], and stimulation of FFAR4 also increases FGF21 expression in adipose tissue via the p38 MAPK signaling pathway [36]. In our experiments, the expression of β3AR and FFAR4 in mice fed the LD and the FOD increased only in BAT and not in inguinal WAT, which may be the reason why FGF21 expression increased only in BAT and not in WAT.

Blood FGF21 levels were lower in mice fed the FOD than in mice fed the LD, unlike the results of FGF21 mRNA expression seen in BAT. It is known that blood FGF21 is mainly derived from liver production [37,38,39] and that adipocyte-derived FGF21 acts locally in an autocrine/paracrine manner [12,40]. Therefore, it is likely that the induction of FGF21 in BAT had no effect on blood FGF21 levels.

In our study, blood FGF21 levels in mice fed the FOD were lower than those in mice fed the LD, and βklotho mRNA expression in the BAT was increased compared to that in mice fed the LD, which may have resulted from an increased sensitivity to FGF21 in the BAT. Previous reports have shown that the expression of βKlotho, a coreceptor of FGF21, is decreased in the obese state [41,42] and that the blood FGF21 concentration is increased due to FGF21 resistance [43,44]. Although we have not found any reports mentioning FO-induced FGF21 sensitivity in adipose tissue, there are suggestions of increased sensitivity to FGF21 with increased expression of βKlotho in the liver by FO [42,45].

Thus, it is possible that induction of FGF21 and increased sensitivity to FGF21 may occur in BAT by feeding FO to DIO mice, but the details and significance of this phenomenon should be verified by future studies.

In conclusion, we have shown that FO increases rectal temperature and exerts obesity-suppressive effects in DIO mice and that FO intake increases the expression of heat-producing genes in BAT but not in WAT.

## Acknowledgement

We wish to thank the Joint Research Laboratory, Kagoshima University Graduate School of Medical and Dental Sciences, for the use of their facilities.

## References

1. Afshin A, Forouzanfar MH, Reitsma MB, Sur P, Estep K, Lee A, et al. Health Effects of Overweight and Obesity in 195 Countries over 25 Years. N Engl J Med. 2017 Jul 6;377(1):13–27. doi: 10.1056/NEJMoa1614362.

2. Global Obesity Observatory. World Obesity Atlas 2023. Available from: https://data.worldobesity.org/publications/?cat=19 Accessed July 31, 2023.

3. Pahlavani M, Razafimanjato F, Ramalingam L, Kalupahana NS, Moussa H, Scoggin S, et al. Eicosapentaenoic acid regulates brown adipose tissue metabolism in high-fat-fed mice and in clonal brown adipocytes. J Nutr Biochem. 2017 Jan;39:101–109. doi: 10.1016/j.jnutbio.2016.08.012.

4. Bargut TC, Silva-e-Silva AC, Souza-Mello V, Mandarim-de-Lacerda CA, Aguila MB. Mice fed fish oil diet and upregulation of brown adipose tissue thermogenic markers. Eur J Nutr. 2016 Feb;55(1):159–69. doi: 10.1007/s00394-015-0834-0.

5. Ishibashi J, Seale P. Medicine. Beige can be slimming. Science. 2010 May 28;328(5982):1113–4. doi: 10.1126/science.

6. Seale P, Bjork B, Yang W, Kajimura S, Chin S, Kuang S, et al. PRDM16 controls a brown fat/skeletal muscle switch. Nature. 2008 Aug 21;454(7207):961–7. doi: 10.1038/nature07182.

7. Wu J, Boström P, Sparks LM, Ye L, Choi JH, Giang AH, et al. Beige adipocytes are a distinct type of thermogenic fat cell in mouse and human. Cell. 2012 Jul 20;150(2):366–76. doi: 10.1016/j.cell.2012.05.016.

8. Kim M, Goto T, Yu R, Uchida K, Tominaga M, Kano Y, et al. Fish oil intake induces UCP1 upregulation in brown and white adipose tissue via the sympathetic nervous system. Sci Rep. 2015 Dec 17;5:18013. doi: 10.1038/srep18013.

9. Yamazaki T, Li D, Ikaga R. Fish Oil Increases Diet-Induced Thermogenesis in Mice. Mar Drugs. 2021 May 17;19(5):278. doi: 10.3390/md19050278.

10. Okue S, Ishikawa E, Nakahara R, Ito T, Okura T, Sakae M, et al. Fish oil suppresses obesity more potently in lean mice than in diet-induced obese mice but ameliorates steatosis in such obese mice. Biosci Biotechnol Biochem. 2021 Feb 18;85(2):421–429. doi: 10.1093/bbb/zbaa038.

11. Chau MD, Gao J, Yang Q, Wu Z, Gromada J. Fibroblast growth factor 21 regulates energy metabolism by activating the AMPK-SIRT1-PGC-1alpha pathway. Proc Natl Acad Sci U S A. 2010 Jul 13;107(28):12553–8. doi: 10.1073/pnas.1006962107.

12. Fisher FM, Kleiner S, Douris N, Fox EC, Mepani RJ, Verdeguer F, et al. FGF21 regulates PGC-1α and browning of white adipose tissues in adaptive thermogenesis. Genes Dev. 2012 Feb 1;26(3):271–81. doi: 10.1101/gad.177857.111.

13. Huang XF, Xin X, McLennan P, Storlien L. Role of fat amount and type in ameliorating diet-induced obesity: insights at the level of hypothalamic arcuate nucleus leptin receptor, neuropeptide Y and proopiomelanocortin mRNA expression. Diabetes Obes Metab. 2004 Jan;6(1):35–44. doi: 10.1111/j.1463-1326.2004.00312.x.

14. De Craemer D, Vamecq J, Roels F, Vallée L, Pauwels M, Van den Branden C. Peroxisomes in liver, heart, and kidney of mice fed a commercial fish oil preparation: original data and review on peroxisomal changes induced by high-fat diets. J Lipid Res. 1994 Jul;35(7):1241–50. Retrieved from https://www.sciencedirect.com/science/article/pii/S0022227520399673?via%3Dihub.

15. Zhang G, Sun Q, Liu C. Influencing Factors of Thermogenic Adipose Tissue Activity. Front Physiol. 2016 Feb 5;7:29. doi: 10.3389/fphys.2016.00029.

16. Seale P, Kajimura S, Yang W, Chin S, Rohas LM, Uldry M, et al. Transcriptional control of brown fat determination by PRDM16. Cell Metab. 2007 Jul;6(1):38–54. doi: 10.1016/j.cmet.2007.06.001.

17. Tseng YH, Kokkotou E, Schulz TJ, Huang TL, Winnay JN, Taniguchi CM, et al. New role of bone morphogenetic protein 7 in brown adipogenesis and energy expenditure. Nature. 2008 Aug 21;454(7207):1000–4. doi: 10.1038/nature07221.

18. Hondares E, Rosell M, Díaz-Delfín J, Olmos Y, Monsalve M, Iglesias R, et al. Peroxisome proliferator-activated receptor α (PPARα) induces PPARγ coactivator 1α (PGC-1α) gene expression and contributes to thermogenic activation of brown fat: involvement of PRDM16. J Biol Chem. 2011 Dec 16;286(50):43112–22. doi: 10.1074/jbc.M111.252775.

19. Enriori PJ, Sinnayah P, Simonds SE, Garcia Rudaz C, Cowley MA. Leptin action in the dorsomedial hypothalamus increases sympathetic tone to brown adipose tissue in spite of systemic leptin resistance. J Neurosci. 2011 Aug 24;31(34):12189–97. doi: 10.1523/JNEUROSCI.2336-11.2011.

20. Izquierdo AG, Crujeiras AB, Casanueva FF, Carreira MC. Leptin, Obesity, and Leptin Resistance: Where Are We 25 Years Later? Nutrients. 2019 Nov 8;11(11):2704. doi: 10.3390/nu11112704.

21. Simonds SE, Pryor JT, Ravussin E, Greenway FL, Dileone R, Allen AM, et al. Leptin mediates the increase in blood pressure associated with obesity. Cell. 2014 Dec 4;159(6):1404–16. doi: 10.1016/j.cell.2014.10.058.

22. Cannon B, Nedergaard J. Brown adipose tissue: function and physiological significance. Physiol Rev. 2004 Jan;84(1):277–359. doi: 10.1152/physrev.00015.2003.

23. Bhaskaran S, Unnikrishnan A, Ranjit R, Qaisar R, Pharaoh G, Matyi S, et al. A fish oil diet induces mitochondrial uncoupling and mitochondrial unfolded protein response in epididymal white adipose tissue of mice. Free Radic Biol Med. 2017 Jul;108:704–714. doi: 10.1016/j.freeradbiomed.2017.04.028.

24. Sadurskis A, Dicker A, Cannon B, Nedergaard J. Polyunsaturated fatty acids recruit brown adipose tissue: increased UCP content and NST capacity. Am J Physiol. 1995 Aug;269(2 Pt 1):E351–60. doi: 10.1152/ajpendo.1995.269.2.E351.

25. Collins S, Daniel KW, Petro AE, Surwit RS. Strain-specific response to beta 3-adrenergic receptor agonist treatment of diet-induced obesity in mice. Endocrinology. 1997 Jan;138(1):405–13. doi: 10.1210/endo.138.1.4829.

26. Fromme T, Klingenspor M. Uncoupling protein 1 expression and high-fat diets. Am J Physiol Regul Integr Comp Physiol. 2011 Jan;300(1):R1–8. doi: 10.1152/ajpregu.00411.2010.

27. Shin W, Okamatsu-Ogura Y, Matsuoka S, Tsubota A, Kimura K. Impaired adrenergic agonist-dependent beige adipocyte induction in obese mice. J Vet Med Sci. 2019 Jun 6;81(6):799–807. doi: 10.1292/jvms.19-0070.

28. Yoneshiro T, Aita S, Matsushita M, Okamatsu-Ogura Y, Kameya T, Kawai Y, et al. Age-related decrease in cold-activated brown adipose tissue and accumulation of body fat in healthy humans. Obesity (Silver Spring). 2011 Sep;19(9):1755–60. doi: 10.1038/oby.2011.125.

29. Du S, Jin J, Fang W, Su Q. Does Fish Oil Have an Anti-Obesity Effect in Overweight/Obese Adults? A Meta-Analysis of Randomized Controlled Trials. PLoS One. 2015 Nov 16;10(11):e0142652. doi: 10.1371/journal.pone.0142652.

30. Zhang YY, Liu W, Zhao TY, Tian HM. Efficacy of Omega-3 Polyunsaturated Fatty Acids Supplementation in Managing Overweight and Obesity: A Meta-Analysis of Randomized Clinical Trials. J Nutr Health Aging. 2017;21(2):187–192. doi: 10.1007/s12603-016-0755-5.

31. Keshavarz SA, Mostafavi SA, Akhondzadeh S, Mohammadi MR, Hosseini S, Eshraghian MR, et al. Omega-3 supplementation effects on body weight and depression among dieter women with co-morbidity of depression and obesity compared with the placebo: A randomized clinical trial. Clin Nutr ESPEN. 2018 Jun;25:37–43. doi: 10.1016/j.clnesp.2018.03.001.

32. Thorsdottir I, Tomasson H, Gunnarsdottir I, Gisladottir E, Kiely M, Parra MD, et al. Randomized trial of weight-loss-diets for young adults varying in fish and fish oil content. Int J Obes (Lond). 2007 Oct;31(10):1560–6. doi: 10.1038/sj.ijo.0803643.

33. Hondares E, Iglesias R, Giralt A, Gonzalez FJ, Giralt M, Mampel T, Villarroya F. Thermogenic activation induces FGF21 expression and release in brown adipose tissue. J Biol Chem. 2011 Apr 15;286(15):12983–90. doi: 10.1074/jbc.M110.215889.

34. Chartoumpekis DV, Habeos IG, Ziros PG, Psyrogiannis AI, Kyriazopoulou VE, Papavassiliou AG. Brown adipose tissue responds to cold and adrenergic stimulation by induction of FGF21. Mol Med. 2011;17(7-8):736–40. doi: 10.2119/molmed.2011.00075.

35. Al Mahri S, Malik SS, Al Ibrahim M, Haji E, Dairi G, Mohammad S. Free Fatty Acid Receptors (FFARs) in Adipose: Physiological Role and Therapeutic Outlook. Cells. 2022 Feb 21;11(4):750. doi: 10.3390/cells11040750.

36. Quesada-López T, Cereijo R, Turatsinze JV, Planavila A, Cairó M, Gavaldà-Navarro A, et al. The lipid sensor GPR120 promotes brown fat activation and FGF21 release from adipocytes. Nat Commun. 2016 Nov 17;7:13479. doi: 10.1038/ncomms13479.

37. Markan KR, Naber MC, Ameka MK, Anderegg MD, Mangelsdorf DJ, Kliewer SA, et al. Circulating FGF21 is liver derived and enhances glucose uptake during refeeding and overfeeding. Diabetes. 2014 Dec;63(12):4057–63. doi: 10.2337/db14-0595.

38. Nishimura T, Nakatake Y, Konishi M, Itoh N. Identification of a novel FGF, FGF-21, preferentially expressed in the liver. Biochim Biophys Acta. 2000 Jun 21;1492(1):203–6. doi: 10.1016/s0167-4781(00)00067-1.

39. BonDurant LD, Potthoff MJ. Fibroblast Growth Factor 21: A Versatile Regulator of Metabolic Homeostasis. Annu Rev Nutr. 2018 Aug 21;38:173–196. doi: 10.1146/annurev-nutr-071816-064800.

40. Justesen S, Haugegaard KV, Hansen JB, Hansen HS, Andersen B. The autocrine role of FGF21 in cultured adipocytes. Biochem J. 2020 Jul 17;477(13):2477–2487. doi: 10.1042/BCJ20200220.

41. Gallego-Escuredo JM, Gómez-Ambrosi J, Catalan V, Domingo P, Giralt M, Frühbeck G, et al. Opposite alterations in FGF21 and FGF19 levels and disturbed expression of the receptor machinery for endocrine FGFs in obese patients. Int J Obes (Lond). 2015 Jan;39(1):121–9. doi: 10.1038/ijo.2014.76.

42. Martínez-Fernández L, González-Muniesa P, Sáinz N, Laiglesia LM, Escoté X, Martínez JA, et al. Maresin 1 Regulates Hepatic FGF21 in Diet-Induced Obese Mice and in Cultured Hepatocytes. Mol Nutr Food Res. 2019 Dec;63(24):e1900358. doi: 10.1002/mnfr.201900358.

43. Fisher FM, Chui PC, Antonellis PJ, Bina HA, Kharitonenkov A, Flier JS, Maratos-Flier E. Obesity is a fibroblast growth factor 21 (FGF21)-resistant state. Diabetes. 2010 Nov;59(11):2781–9. doi: 10.2337/db10-0193. Epub 2010 Aug 3.

44. Tan BK, Hallschmid M, Adya R, Kern W, Lehnert H, Randeva HS. Fibroblast growth factor 21 (FGF21) in human cerebrospinal fluid: relationship with plasma FGF21 and body adiposity. Diabetes. 2011 Nov;60(11):2758–62. doi: 10.2337/db11-0672.

45. Yang W, Chen X, Liu Y, Chen M, Jiang X, Shen T, et al. N-3 polyunsaturated fatty acids increase hepatic fibroblast growth factor 21 sensitivity via a PPAR-γ-β-klotho pathway. Mol Nutr Food Res. 2017 Sep;61(9). doi: 10.1002/mnfr.201601075.

